# Safety and Feasibility of Infusing *Ex Vivo* Expanded Allogeneic Canine Natural Killer Cells for the Treatment of Metastatic Solid Tumors

**DOI:** 10.64898/2026.03.19.712729

**Authors:** Allison M. Weisnicht, Fernanda Szewc, Monica Moonkyung Cho, Han-Yun Hannah Cheng, Swetha Ganesh, Lauren Mahoney, Kathryn Fox, Portia R. Smith, Mallery Olsen, Rebecca M. Richards, David M. Vail, Christian M Capitini

## Abstract

**Background:** Companion canines need advances in therapeutic options for solid tumor malignancies. Prior studies established feasibility of autologous natural killer (NK) cell infusions in canines with solid tumors; however, autologous products are limited by dysfunctional immunity and a manufacturing process that delays care. Allogeneic NK cells offer the possibility of “off-the-shelf” therapy to be administered from healthy donors.

**Methods:** Peripheral blood mononuclear cells (PBMCs) were isolated from healthy canine donors via density gradient separation. NK cells were expanded with recombinant human IL-2 and canine IL-21 with the addition of K562 feeder cells transfected with CD137 ligand and membrane bound human IL-15. Additional experiments included IL-12 in the expansions. In vitro potency was assessed via co-culture with the D17-mKate2 canine osteosarcoma cell line. Three canines were enrolled in a phase 1 trial infusing ex vivo expanded allogeneic NK cells after lymphodepletion.

**Results:** Flow cytometric analysis confirmed successful expansion of canine NK cells with up to 50% of cells demonstrating NKp46+ after 14 days of expansion. Residual T cell numbers varied based on donor. The addition of IL-12 led to increased NK cell expansion. Incucyte demonstrated potency with increasing osteosarcoma cell death at higher effector to target ratios. Three canines with metastatic/refractory solid tumors were successfully lymphodepleted and infused with allogeneic NK cell products. The canines tolerated the infusions well.

**Conclusions:** Canine allogeneic NK cells were successfully expanded and activated ex vivo, demonstrated potency in vitro, and safety in vivo. Further studies will optimize the NK cell product and escalate dosing to reach the maximal tolerable dose.

## Introduction

Companion canines develop a similar set of malignancies as humans, including melanoma, lymphoma, and osteosarcoma. In fact, canines spontaneously develop osteosarcoma at an estimated rate that is 27-fold higher than humans.^1^ Canine osteosarcoma is also known to have genetic and phenotypic similarities to human osteosarcoma.^1^ Compared to laboratory mouse models, companion canines provide a unique heterogenous population that retains the host immune system and tumor microenvironment. Osteosarcoma is the most common bone malignancy in children. Treatment of osteosarcoma has not significantly changed over the past 50 years. The outcomes for metastatic osteosarcoma remain dismal, with a 5-year overall survival around 30%.^2–4^ Outcomes are even worse for patients who have relapsed. Novel therapies with decreased toxicities are required to improve outcomes as well as the quality of life for children and companion canines. Cellular therapies, and in particular natural killer (NK) cells, are a promising therapy for the treatment of osteosarcoma.^5^

NK cells are part of the innate immune system with the capacity to kill cells that are damaged, virally infected, or malignantly transformed without a need for prior exposure.^6^ NK cells are also able to recognize and subsequently kill their target cells without engaging the major histocompatibility complex (MHC) and in a tumor-agnostic manner.^7^ These features of NK cells position them as a compelling choice in the development of a cellular therapy product for the treatment of high-risk malignancies. As compared to adoptive cellular therapy with T-cells that require the product to be autologous due to inherent risk of graft-versus-host disease (GVHD) when using an allogeneic product, NK cells do not carry this same risk and can be used “off the shelf” to treat patients with cancer in a more expedited fashion.^8^ For example, manufacturing of autologous chimeric antigen receptor T (CAR T) cells is at least a three-week process that involves apheresis of T cells from the patient, *ex vivo* modification of those T cells typically through retro- or lentiviral transduction of an antigen-specific receptor, and then reinfusion of the CAR T cells into the patient.^9^ Because patients are heavily pre-treated, autologous T cells can be suboptimal for generating maximal anti-tumor cytotoxicity. Allogeneic NK cells are an attractive alternative cellular therapy since they can be generated from healthy donors. There is also interest in how NK cells could be used in combination with conventional therapies like standard chemotherapy and radiation regimens as well as other immunotherapy such as immune checkpoint inhibitors and monoclonal antibodies.^10^

Prior to this study, there have been three clinical trials studying canine NK cell therapy. Canter et al. performed intratumoral injections of autologous *ex vivo* expanded NK cells post-radiation therapy in canines with osteosarcoma. They reported increased cytotoxicity and NK cell homing with prior radiation exposure.^11^ Five canines remained free of metastasis at the 6-month primary endpoint and one canine had resolution of suspicious pulmonary nodules. The same group then studied both autologous and allogeneic peripheral blood mononuclear cell (PBMC)-expanded NK cells in canine patients with osteosarcoma and melanoma.^12^ They demonstrated safety and feasibility of using unmanipulated PBMCs (no T cell depletion step) in autologous (n=9) and allogeneic (n=5) NK cell infusions.

This pilot study focuses on the production, safety, and feasibility of canine allogeneic NK cell infusions. An allogeneic approach was chosen due to multiple benefits over the use of an autologous product from both biological and feasibility standpoints. Biological factors include the use of healthy NK cells without prior exposure to chemotherapy or other treatments, overcoming tumor escape mechanisms from the host immune system (including inhibition by self MHC molecules), and results from autologous canine NK infusions that showed limited treatment beneft.^12^ The results from 4 allogeneic *ex vivo* expanded NK cell infusions (1.5 – 5 x 10^5^ NK cells/kg) given to 3 canine subjects with metastatic cancer are reported.

## Methods

### Canine Peripheral Blood Mononuclear Cell (PBMC) isolation

Fifty milliliters of whole blood was collected from healthy canine donors at the University of Wisconsin School of Veterinary Medicine (Table 1). One milliliter of 1000 units/mL of heparin was added for each 50ml of blood. Whole blood was stored at room temperature if PBMC isolation was completed on the same day as collection, otherwise it was stored at 4 degrees Celsius for up to 48 hours prior to PBMC isolation. Sterile phosphate-buffered saline (Corning, catalog#MT21040CV) was added 1:1 to the whole blood sample. PBMCs were isolated from whole blood using sterile Ficoll media for density gradient centrifugation (Ficoll-Paque, Cytiva, catalog#17144003). Red blood cell lysis was completed using ammonium-chloride-potassium (ACK) lysing buffer (Thermo Scientific Chemicals, catalog#J62990.AK). Isolated PBMCs were then suspended in media at 1×10^6^ cells/mL. Media used for all cell cultures consisted of RPMI-1640 (Corning, catalog#10-040-CV), 10% fetal bovine serum (Foundation, catalog#900-108), and L-glutamine (Corning, catalog#25-005-CI) with the addition of 100 units/ml penicillin, 100 μg/ml streptomycin, and 2.5 mg/ml Plasmocin (Plasmocin, catalog#ant-mpp) as antibiotic prophylaxis.

**Table 1.** Characteristics of canine whole blood donors.

| Canine Donor | Age (years) | Breed | Weight (kg) | Sex | PBMC Count (x 10<sup>6</sup>) |
| --- | --- | --- | --- | --- | --- |
| 1 | 5 | Treeing walking coonhound | 30.2 | Male, neutered | 168.6 |
| 2 | 2 | Mixed hound | 48.8 | Male, neutered | 124.8 |
| 3 | 4 | Standard poodle | 14.5 | Female, spayed | 71.8 |
| 4 | 11 | Standard poodle | 19.4 | Female, spayed | 71.4 |
| 5 | 2 | Golden retriever | 30.0 | Female, spayed | 34.0 |
| 6 | 8 | Mix | 14.5 | Female, spayed | 65.5 |

### T cell depletion

T cell depletion was performed on PBMCs on day 0 to enrich for NK cells and on day 14 of expansion to reduce risk of GVHD from any potential T cell contamination. CD3 positive T cells were first labeled with a FITC conjugated antibody (Bio Rad antibodies, catalog#MCA1774F). CD3 depletion was then completed with EasySep FITC Positive Selection Kit (Stemcell Technologies, catalog#17682) via magnetic cell separation. The negative cell fraction was collected and placed in culture media at 37°C at 5% CO_2_.

### Canine NK cell expansion

After isolation, T cell depleted PBMCs underwent expansion using feeder cells and cytokines (Figure S1). Briefly, 100 IU/mL recombinant human interleukin-2 (rhIL2) (Biological Resources Branch, National Cancer Institute-Frederick) and 5ng/mL canine IL-21 (Biotechne, catalog#5849-ML) were added with fresh media every 2-3 days to the cell culture. Five ng/mL canine IL-12 (Biotechne, catalog#2118-CL) was also added to some expansions. Human K562 feeder cells transfected with 4-1BB ligand and membrane bound (mb) IL15 (K562-41BBL-mbIL15), a gift from Dario Campana (National University of Singapore), were irradiated at 100 gray (Gy) using a cell irradiator (Xstrahl CIX2) and cultured in media at 37°C at 5% CO_2_. Irradiated feeder cells were added to NK cells on days 0 and 7 of the expansion. Cells were counted via a hemocytometer to enumerate expansion. Flow cytometry was completed around days 7 and 14 during the expansion to specifically monitor NK cell and T cell population percentages.

### Flow cytometry

Cells were incubated with Fc receptor blocking solution (canine Fc Receptor Binding Inhibitor Polyclonal Antibody, Invitrogen, Catalog#14-9162-42) and subsequently stained with fluorophore-conjugated antibodies. The following antibodies were used in analyses: mouse anti-canine CD3:FITC (clone CA17.2A12, BioRad Catalog#MCA1774F), mouse anti-canine unconjugated NKp46 (CD335) (clone 48A, Millipore Sigma, catalog#MABF2109-100UG) that was secondarily detected with anti-mouse APC (clone RMG2a-62, BioLegend, catalog#407110), rat anti-canine CD8a: PerCP-eFluor 710 (clone YCATe55.9, Invitrogen, catalog#46-5080-42), rat anti-canine CD4:Star Bright Violet 610 (clone YKIX302.9, BioRad, catalog#MCA1038SBV610) and mouse anti-human IL15:PE (clone 34559, Biotechne, catalog#IC2471P). Anti-canine NKp46 was used to define the canine NK cell population and anti-canine CD3, CD4, and CD8 were used to define the canine T cell population. Elimination of feeder cells was confirmed via flow cytometry analysis using anti-human IL15, which is expressed on the surface of the K562-41BBL-mbIL15 feeder cell line. All flow cytometry studies were conducted on an Attune 4-laser, 14-color analysis cytometer (Thermo Fisher Scientific, Waltham, MA, USA). FlowJo Software was used for flow analyses (FlowJo Software v10.10, Ashland, OR, USA).

### Incucyte

The Incuyte live cell imaging system (Sartorius, Göttingen, Germany) was used to analyze cytotoxicity of the *ex vivo* expanded canine NK cell product *in vitro*. D-17 (ATCC, Manassas, Virginia), previously derived from a poodle with osteosarcoma, was transfected with an mKate2 reporter gene (D17-mKate2) and cultured in media at 37°C at 5% CO_2_. ATCC guidelines were followed for cell authentication of all cell lines using morphology monitoring, growth curve analysis, and testing for mycoplasma within 6 months of use. D17-mKate2 cells were cultured in media and plated at 5,000 cells per well in a 96-well plate. Effector cells were plated at effector:target (E:T) ratios of 1:1, 2:1, and 5:1. Images were taken every 4 hours for 72 hours to assess red object intensity over time in various co-culture conditions. A positive control included addition of 3μM staurosporine per D17-Kate2 well. Data analyses were completed using GraphPad Prism version 10 (GraphPad Software, LLC, San Diego, CA, USA).

### Release criteria for fresh allogeneic NK cell products

The use of companion animals in our pilot study was approved by the Institutional Animal Care and Use Committee (IACUC) at the University of Wisconsin-Madison. Briefly, peripheral blood was collected from healthy canine donors and NK cells were isolated from T cell depleted PBMCs, expanded and activated as above. Peripheral blood collections were performed from each canine donor with the goal of generating enough cells for one intravenous (IV) NK infusion at 1 x 10^5^ NK cells/kg as a fresh infusion. Mycoplasma testing was performed on each cell product prior to infusion via MycoStrip Mycoplasma Detection Kit (InvivoGen, catalog#rep-mys-10). Endotoxin testing, to ensure sterility of the products, was performed via the Limulus Amebocyte Lysate (LAL) test in the University of Wisconsin Program for Advanced Cell Therapy (PACT) lab. Release criteria for each infusion product included viability greater than or equal to 50%, 5×10^6^ NK cells per kilogram +/- 20% (using recipient’s weight), less than 1×10^5^ T cells per kilogram (using recipient’s weight), negative for mycoplasma, and endotoxin less than 5 EU/mL.

### Lymphodepletion and adoptive transfer of allogeneic NK immunotherapy in canines with metastatic refractory malignancies

Baseline assessments included history, physical exam, verification of diagnosis of metastatic cancer, tumor imaging by radiography, ultrasonography, or computed tomography, a complete blood count (CBC), serum or plasma biochemical profile, and urinalysis. Eligible canine subjects underwent lymphodepletion with 400mg/m^2^ oral cyclophosphamide on day -4 and 10mg/m^2^ IV fludarabine on day -3. One dose of 1mg/kg oral furosemide was provided after administration of cyclophosphamide to mitigate bladder toxicity. Just prior to infusion, allogeneic donor NK cells were resuspended in 1mL/kg of sterile 0.9% NaCl and intravenously infused on day 0 in eligible canine patients using a 19G catheter at a rate of 2 mL per 3 minutes. RhIL-2 was co-injected subcutaneously at a dose of 250,000 IU/kg on days 0, +7, and +14. Canines were re-evaluated once weekly post-infusion (Figure 1).

**Figure 1.**
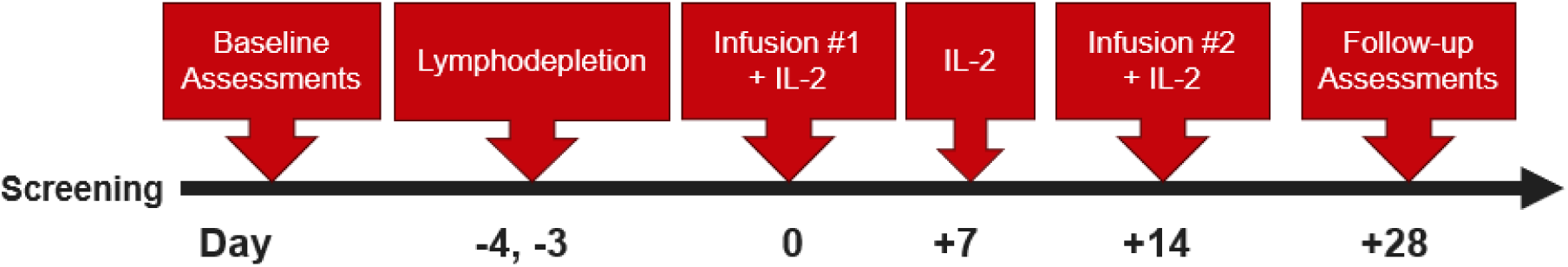
Canine NK cellular therapy trial schematic. Baseline assessments were completed prior to enrollment (history, physical examination, imaging of primary disease site, and labs). Lymphodepletion with cyclophosphamide and fludarabine was completed on days -4 and -3. NK cell infusions were completed on days 0 and +14 with co-infusion of IL-2. IL-2 also administered on day +7 to support NK cell activation *in vivo.* Follow-up assessments were planned for day +28 or at the discretion of the treating veterinarian.

### Monitoring of subjects

Vital signs (i.e., temperature, pulse, respiration, and blood pressure) were monitored hourly for 4 hours post-infusion. Each canine was re-evaluated weekly by a veterinary oncologist for 3 weeks post-infusion with history, physical exam and CBC performed at each visit. Adverse events were graded per VCOG-CTCAE v2.^13^ Follow-up imaging was not required on study since this was a pilot feasibility study, however imaging was allowed as needed for clinical management of new symptoms.

## Results

### Ex vivo expanded NK cells derived from T cell depleted canine PBMCs

First, we wanted to determine the feasibility of expanding allogeneic canine NK cells *ex vivo* using cytokines and feeder cells. Fifty milliliter blood samples were collected from healthy donors of various breeds (Table 1), PBMCs were isolated and depleted of T cells. The cell product was expanded with irradiated K562-41BBL-mbIL15 feeder cells in rhIL-2 and canine IL-21. Cell counts were obtained every few days during expansion (Figure 2A). Flow cytometry was completed on days 0, 7, and 14 for select expansions (Figure 2B) to confirm identity.

**Figure 2.**
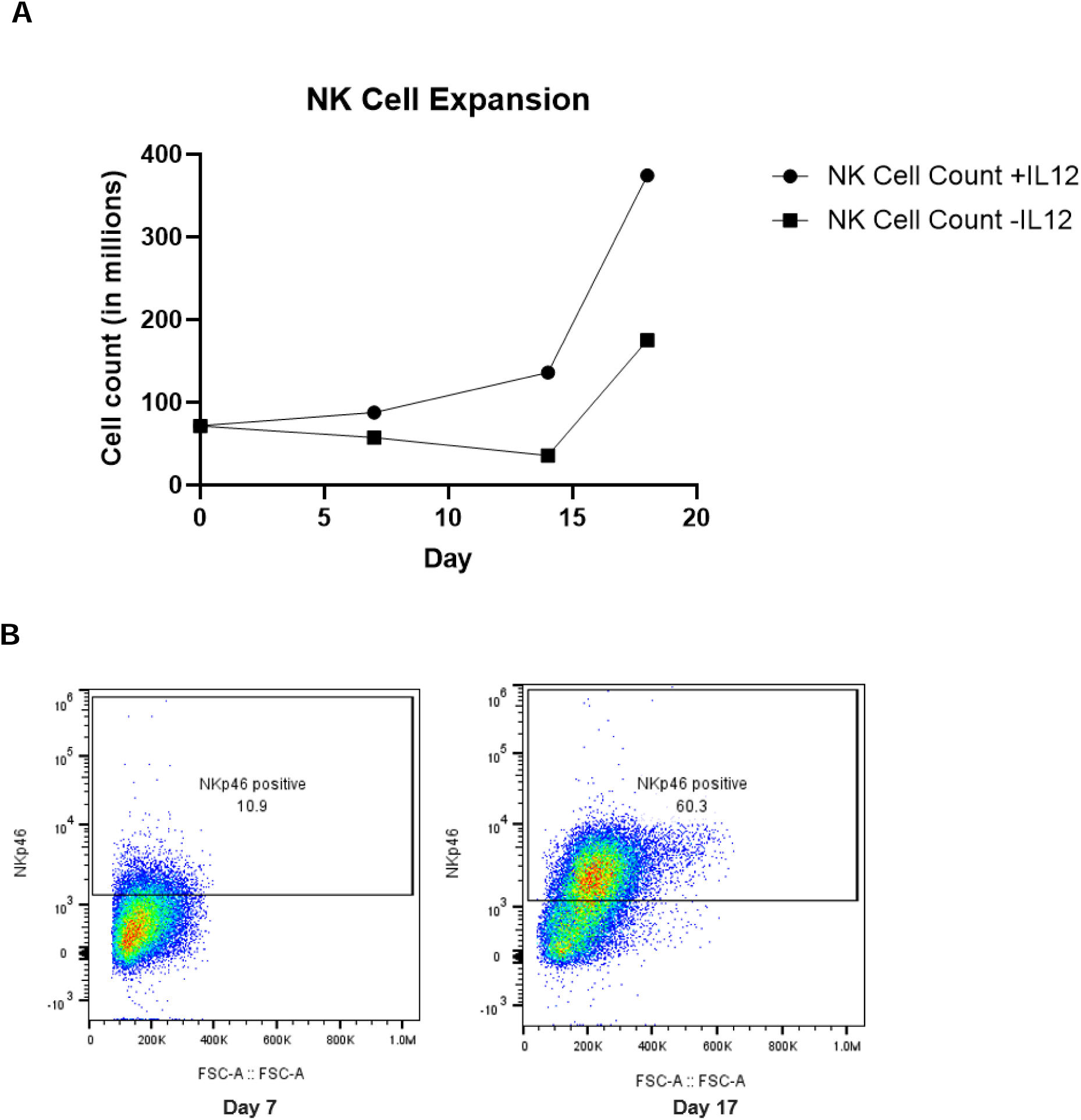
Successful PBMC-derived canine NK cell expansion data. (A) Notable expansion occurred after day 14. The addition of IL-12 resulted in greater NK cell expansion (discussed later). Data shown are representative from 2 expansions. (B) Following the protocol described in Figure S1, the proportion of NK cells of the live co-cultured PBMCs increased from 10.9% to 60.3% as demonstrated by flow cytometry.

### T cell (CD3) depletion

Prior work by others debated the utility of depleting T cells from PBMCs in the setting of providing cytokines and feeder cells that enrich for NK cells.^12^ Thus, CD3 depletion was performed either on day 0 to enrich for NK cells, on day 14 to reduce contaminating T cells, or not at all with four separate donors to assess optimal timing and/or necessity for T cell depletion during *ex vivo* NK cell expansion. CD3 depletion on day 0 resulted in an overall increased NKp46+ population and, in particular, an increased single positive NKp46 population compared to depletion on day 14 and no depletion at all (Figure S2). The single positive NKp46 population is described as a more cytotoxic subset of canine NK cells.^14,15^

### Addition of canine IL-12

Despite demonstrating adequate expansion of NK cells with K562-41BBL-mbIL15 feeder cells, IL-2 and IL-21 (Figure S1), there was inconsistency in successful expansion based on the canine donor. Given these inconsistencies, addition of other cytokines was explored to determine if this would lead to further benefit and increased reliability of NK cell expansion. Based on prior human and murine NK cell studies, the addition of IL-12 was studied to assess for improved canine NK cell expansion.^16,17^ Canine IL-12 was added every 2-3 days in addition to rhIL-2 and canine IL-21 as per the initial protocol. This process was repeated with four separate donors to assess for reproducibility of results. The addition of IL-12 consistently yielded increased canine NK cells as compared to expansions performed without the addition of IL-12 (Figures 2A and 3A). Notably, a CD8^+^NKp46^+^ double positive population appeared to specifically expand with the addition of IL-12 (Figure 3B).

**Figure 3.**
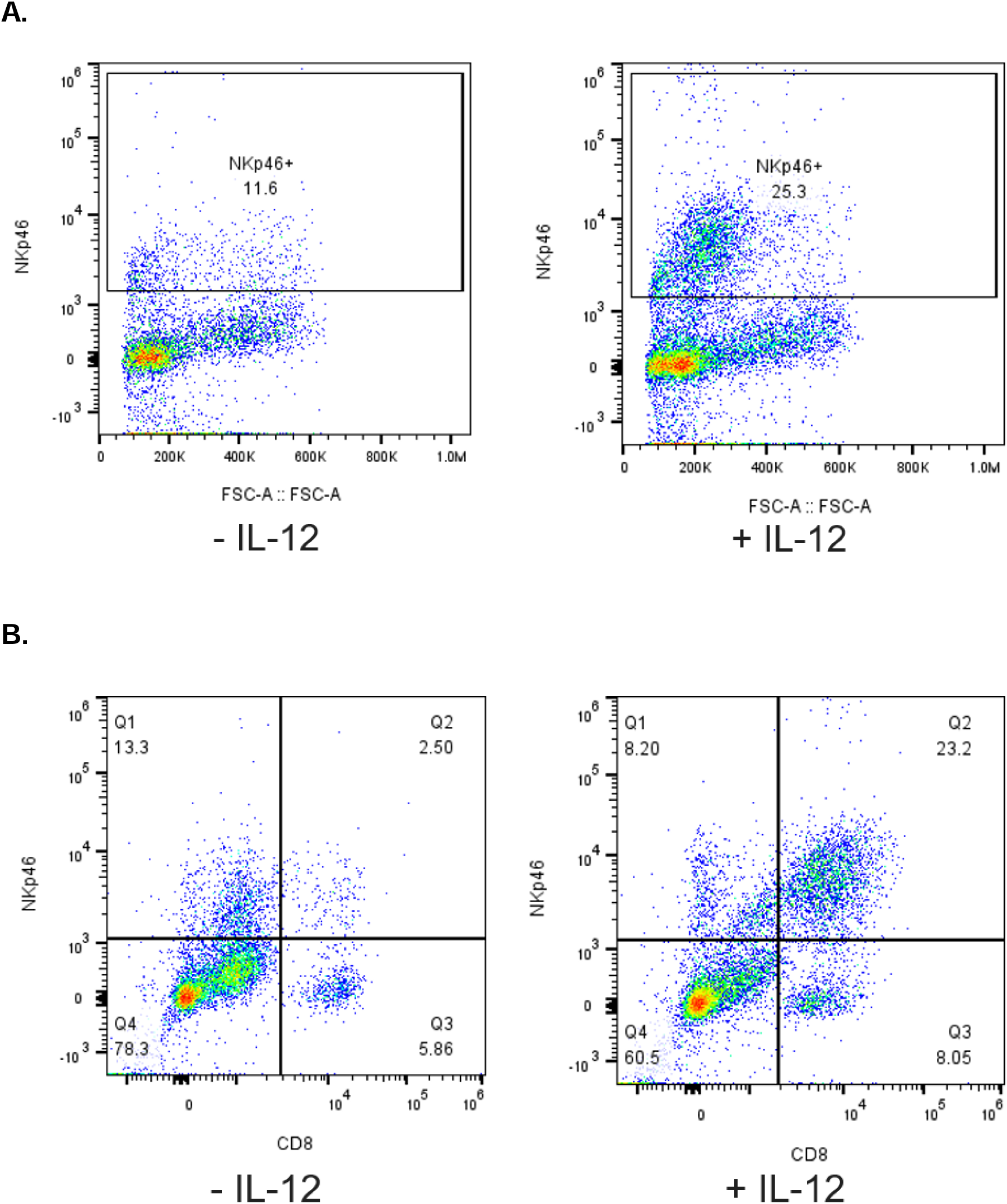
PBMC-derived NK cell expansion with IL-12 leads to an increased NK cell population co-expressing CD8. (A) The addition of IL-12 yielded increased NK cell expansion as shown in the flow cytometry plot. (B) The flow cytometry plots demonstrate expansion of double-positive NKp46+CD8+ population with the addition of IL-12.

### Functional assessment of *ex vivo* expanded NK cells

Incucyte live cell imaging analyses using a co-culture of PBMC-expanded allogeneic NK cells with D17-mKate2 osteosarcoma cells were performed with three separate canine blood donors to assess NK potency. Cytotoxicity was demonstrated with two separate donors without IL-12 at 1:1, 2:1, and 5:1 ratios (Figure 4A and 4B). Flow cytometry analysis was performed just prior to Incucyte set-up to evaluate and correlate the phenotype of effector cells with the cytotoxicity results. Notably, there was significant cytotoxicity seen in an expansion without IL-12 (Figure 4C), however when correlating this data with flow cytometry, the predominant population present were T cells as compared to NK cells (Figure 5). Thus, the addition of IL-12 to the expansion protocol is not anticipated to result in decreased cytotoxicity of NK cells, but rather selectively enriches for NK cells versus T cells.

**Figure 4.**
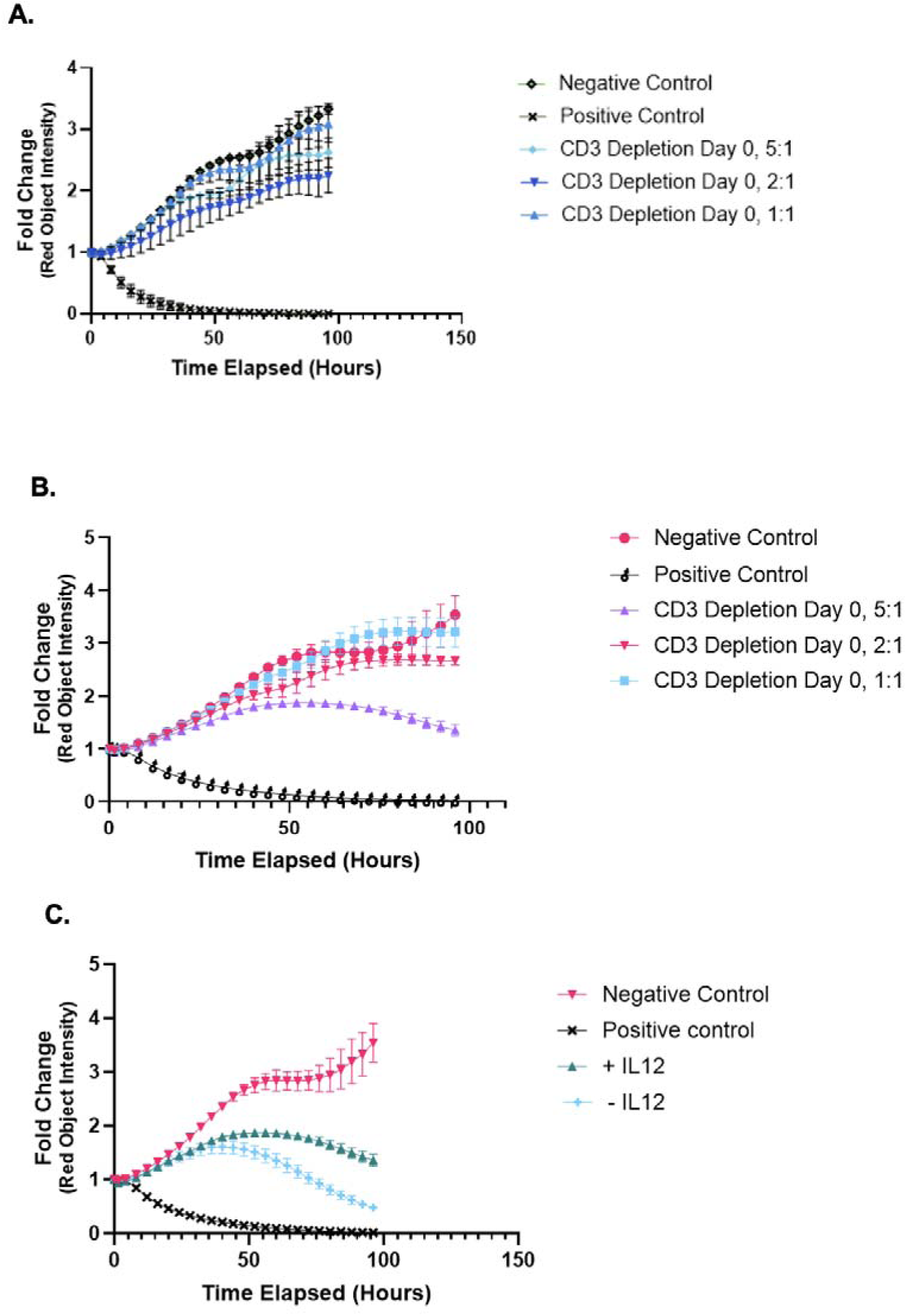
*In vitro* cytotoxicity assays of NK cells co-cultured with a poodle osteosarcoma cell line. (A)+(B) Incucyte live imaging analysis of D17-mKate2 cell line in co-culture with PBMC-derived NK cells at 5:1, 2:1, and 1:1 effector to target ratios from two separate donors. (C) Incucyte live imaging analysis of D17-mKate2 cell line in co-culture with PBMC-derived NK cells with or without IL-12.

**Figure 5.**
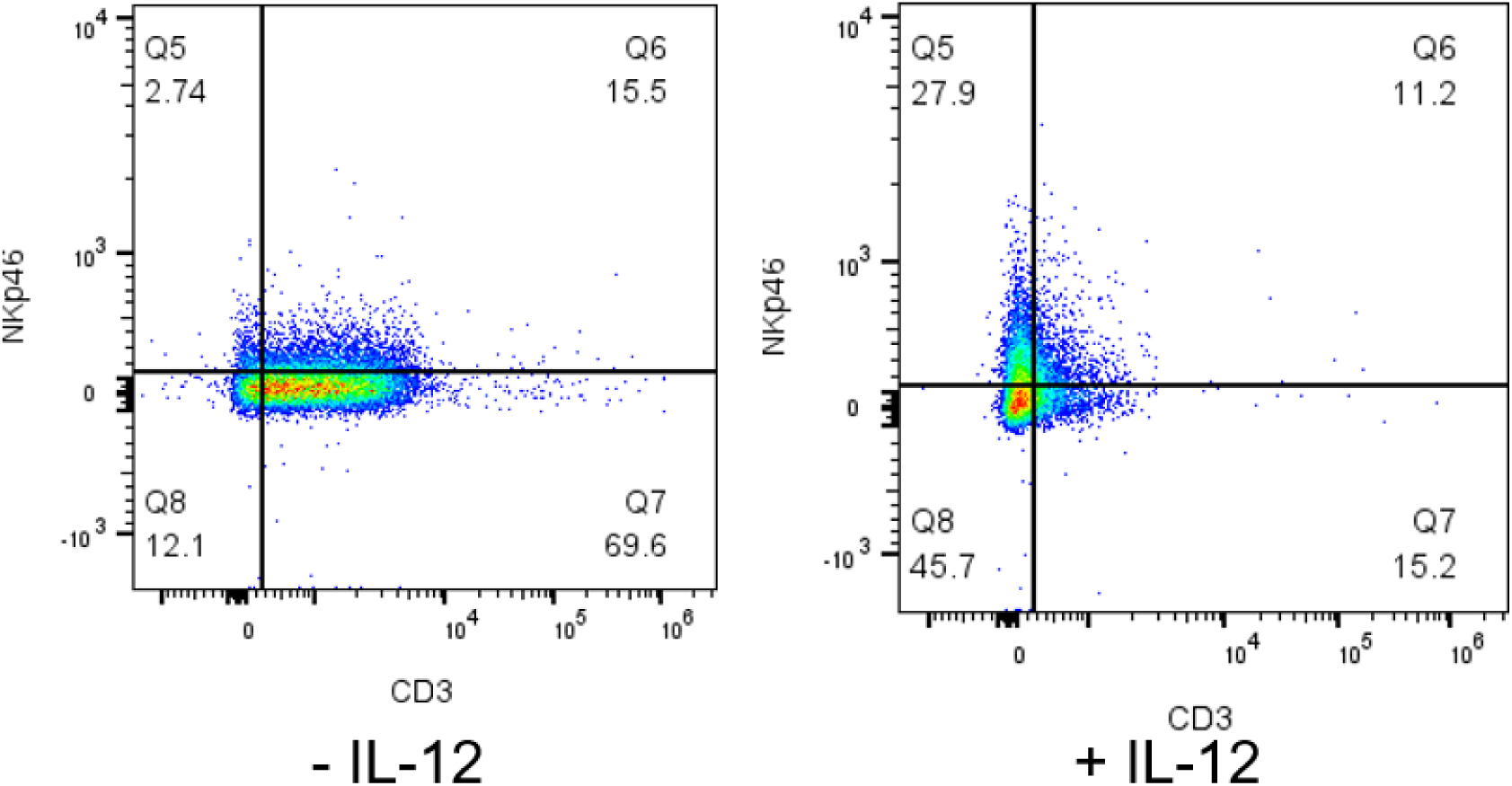
T cell content of NK cell expansions with or without IL-12. Flow cytometry plot demonstrates significant T cell population in expansion without IL12 as compared to NKp46 single positive population with the addition of IL12.

### Phase 1 pilot clinical trial

Four allogeneic *ex vivo* expanded NK cell infusions (1.5 – 5 x 10^5^ NK cells/kg) were safely completed in three canine subjects with metastatic cancer (Table 2). Four subjects were initially screened with 3 subjects eligible to receive infusions. Canine 4 had rapidly progressive disease and was compassionately euthanized before proceeding with enrollment on the trial. All patients tolerated lymphodepletion with cyclophosphamide and fludarabine well, with no reports of bladder toxicity. Lymphocyte reduction was successfully achieved in all 3 subjects, as evidenced by peripheral CBCs (Figure 6). During the NK cell infusion monitoring period, no significant changes were noted, except for a 1^0^F body temperature elevation 2-hours post infusion in one canine.

**Figure 6.**
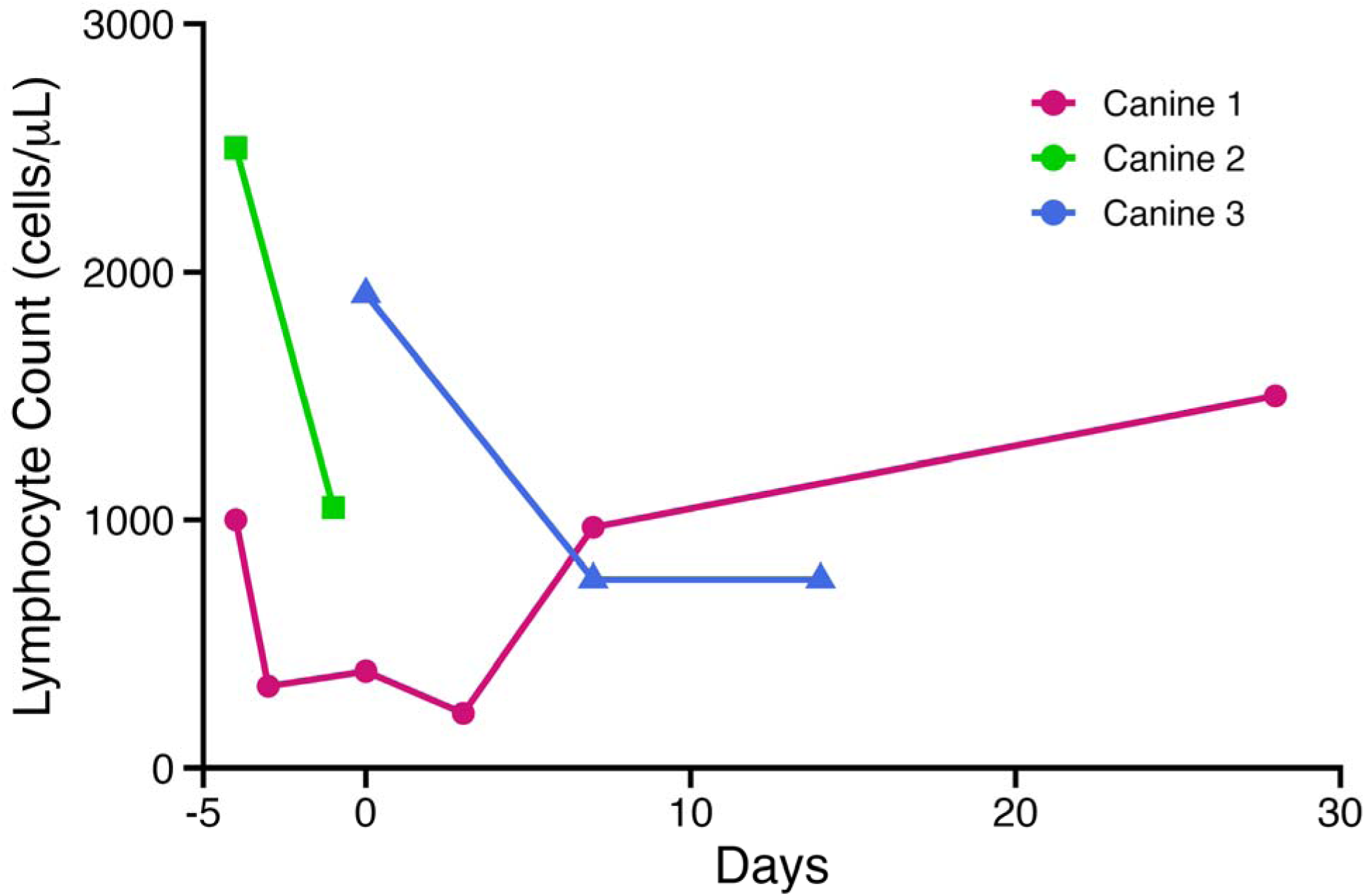
Successful lymphodepletion from fludarabine and cyclophosphamide. Eligible canine subjects underwent lymphodepletion with 400mg/m^2^ oral cyclophosphamide on day -4 and 10mg/m^2^ IV fludarabine on day -3. Allogeneic *ex vivo* expanded canine NK cells were infused on day 0. Lymphocyte depletion demonstrated on peripheral blood CBCs from canines 1, 2, and 3.

**Table 2.**
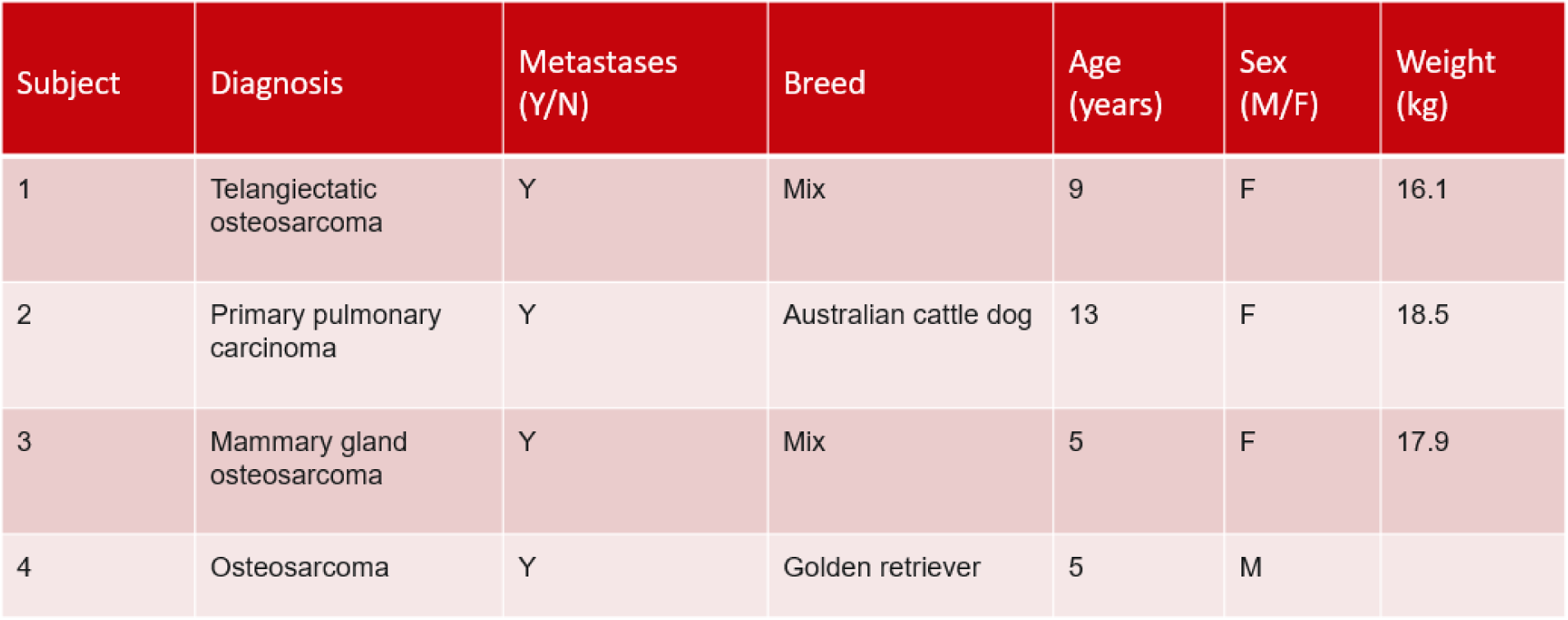
Characteristics of 4 canine patients screened for the allogeneic NK cell infusions.

Patient 1 received 2 infusions one week apart. The doses of *ex vivo* expanded allogeneic NK cells she received were 1.7 x 10^5^ NK cells/kg and 5.0 x 10^5^ NK cells/kg for doses 1 and 2 respectively. The only adverse event was a transient grade 4 episode of fever and neutropenia on day 7, on which she was seen by her local veterinarian for blood cultures (that remained negative) and one dose of IV antibiotic in an ambulatory setting and had a full recovery. She was euthanized on day +57 in the setting of progressive disease.

Patient 2 received one infusion at a dose of 4×10^5^ NK cells/kg and presented one week later with intractable seizures and was subsequently euthanized. Because it was unclear if this was due to the NK cell infusion or not, she underwent necropsy and was found to have extensive brain metastases as the likely etiology of her seizure presentation.

Patient 3 received one infusion at a dose of 1.5×10^5^ NK cells/kg. She was euthanized at day +7 post-infusion in the setting of progressive disease with enlarging malignant (confirmed cytologically) pleural effusions (Figure 7). All allogeneic NK cell infusions contained less than 1×10^5^ T cells to avoid the risk of graft-versus-host disease (GVHD). There was no evidence of GVHD in any of the canines who received the allogeneic NK cell infusions.

**Figure 7.**
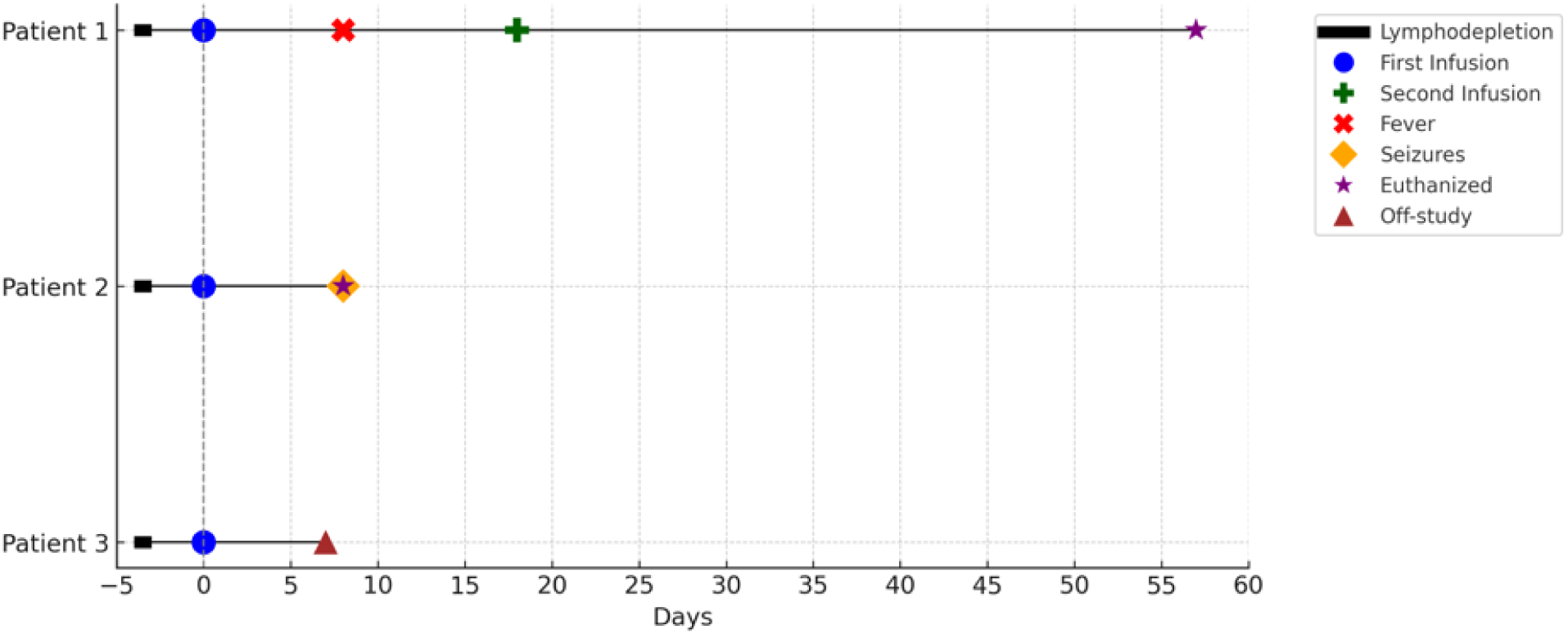
Outcome of canines who received PBMC-derived allogeneic NK cell infusions. The major events of each of the three canine patients are shown in a Swimmer’s plot – main events included lymphodepletion, infusions, fevers, seizures, euthanasia, and off-study time points.

## Discussion

Our studies demonstrated successful *ex vivo* expansion of canine NK cells from a starting population of T cell depleted PBMCs derived from healthy donor peripheral blood samples. Donor-to-donor variability in the final *ex vivo* expanded NK products was observed in terms of resultant NK and T cell population numbers, which is not uncommon in working with dynamic cellular therapy products. We completed further experiments to optimize the quality of the NK product via addition of IL-12 as well as determining the optimal timing for T cell depletion, ultimately completing 4 allogeneic NK cell infusions in 3 companion canines with relapsed/refractory metastatic solid tumors.

One major barrier to canine NK cell research is the lack of consensus on cell surface markers used to identify the NK population.^14^ Some groups identify the canine NK cell population as CD5^dim^, however this has also been found to incompletely capture subsets of cells that have demonstrated NK cell-like activity. For the purposes of our experiments, we defined the canine NK cell population as NKp46^+^CD3^+/-^CD8^+/-^CD4^-^based on general consensus in the field.^9,10,12^ We defined the canine T cell population as NKp46^-^CD3^+^CD8^+/-^CD4^+/-^.^15^ Defining the T cell population is critical in allogeneic cell therapy studies to avoid GVHD. As the canine NK cell research field expands, there needs to be improved uniformity in how researchers define these populations through enhanced classification based on agreed upon phenotype and genotype features. There is also developing work to look at gene expression profiling and transcriptomic analyses of canine NK cells, which could continue to advance the field beyond using surface level phenotypic markers for cell type identification. ^18,19^

The natural dynamic variability that occurs in working with cellular therapy products poses a challenge to manufacturing a consistent product with a reliable workflow. We found that compared to prior published work, using rhIL-2, rhIL-15, and canine IL-21 from bulk-derived PBMCs inconsistently yielded a product that met release criteria for T cell content (<1×10^5^ T cells).^12^ This prompted experimentation with T cell depletion strategies to consistently yield a product with lower T cell content to allow for delivery of a higher dose of NK cells without concern for inducing GVHD. Based on the experiments we performed comparing T cell depletion on days 0 versus 14 versus not at all, T cell depletion on day 0 of expansion was the most consistent in reducing the final T cell population and increasing NK cell expansion, presumably due to decreased T cell competition for cytokines from the start of expansion.

Another strategy employed to assist in NK cell expansion was the addition of IL-12. Use of IL-12 in *ex vivo* expansion of human NK cells has been shown to sustain NK cell survival.^17^ This has additionally been demonstrated in mice.^16^ Given the known molecular homology among mammals, there was interest in studying IL-12 in *ex vivo* canine NK cell expansion. Additionally, a canine-specific IL-12 product is commercially available for research use. As far as we are aware, this is the first described use of IL-12 for *ex vivo* expansion of canine NK cells. We demonstrated that the addition of IL-12 every 2-3 days during NK expansion, in combination with IL-2 and IL-21, yields an increased population of NK cells and particularly enriched for a NKp46^+^CD8a^+^ double positive population. This double positive population in canines has been associated with increased cytotoxicity of tumors.^20^ CD8a is also present on human NK cells, however its presence has been associated with a suppressive effect on NK cell activating receptors.^21^ Thus, further studies are needed to better characterize this NKp46^+^CD8a^+^ double-positive population that is enriched with the addition of IL-12, and determine if it is an NKT cell or an NK cell subset with aberrant expression of CD8a.

Adoptive transfer of autologous NK cells has limited therapeutic benefit due to inhibition by MHC molecules on the tumor, thus we proposed using allogeneic NK cells to enhance alloreactivity against the tumor. Three canines received a total of 4 allogeneic NK cell infusions (1.5 – 5 x 10^5^ NK cells/kg) in combination with rhIL-2 on a phase 1 pilot clinical trial. All three canines tolerated the infusions well with the highest VCOG-CTCAE v2 graded AE as a transient grade 4 event (fever and neutropenia requiring outpatient management). Further studies will dose-escalate towards the intended goal of infusing 5×10^6^ NK cells/kg. The limited sample size and extensive baseline disease of participants at time of trial enrollment preclude the ability to assess for efficacy but allowed for establishment of feasibility and preliminary safety assessment, especially given limited published experience using lymphodepletion with fludarabine in dogs. The canines enrolled on our trial to date had widely metastatic disease at diagnosis thus limiting analyses in post-infusion setting, although future trials could include peripheral blood collections for extensive immunophenotyping and cell-free tumor DNA assessments to assess disease response without imaging.^22,23^

To date, there are only 3 published trials describing the use of *ex vivo* expanded canine NK cells. This report of our 3 canine patients (4 infusions) contributes to the limited but growing work in this area. Progress for the future in this field relies on establishing consistent definitions for identification and characterization of canine NK cell population subsets^18,19^, reliable *ex vivo* expansion NK manufacturing protocols, dose-escalation studies to demonstrate maximal tolerable NK doses, and tracking of NK cells *in vivo*.^24,25^ Additionally, combinatory therapeutic strategies (i.e., NK cellular therapy plus chemotherapy or other immunotherapy agents or CAR NK approaches) could be explored, which could be optimized in canine patients in the future.

## Conclusions

We demonstrated the feasibility and safety of infusing fresh allogeneic *ex vivo* expanded NK cells in companion canines with metastatic solid tumors. While there is great variability in final NK cell and T cell yields from different healthy donors despite similar expansion protocols, the addition of IL-12 and incorporation of T cell depletion resulted in increased NK cell numbers and a product that showed potency *in vitro*. A major limitation of our study includes the small number of patients and low NK cells doses (1.5 – 5 x 10^5^ NK cells/kg). We plan to optimize the NK cell expansion methodology to escalate NK cell dosing to a target infusion dose of 5×10^6^ cells/kg to better demonstrate efficacy with ongoing safety and feasibility. This interim trial report contributes to the development of novel adoptive cell therapies for companion canines with cancer.

## Declaration of interests

CMC reports honorarium from Bayer and Novartis, as well as equity interest in Elephas, for advisory board memberships. These companies had no input in the study design, analysis, manuscript preparation, or decision to submit for publication. No other relevant conflicts of interest are reported.

## Acknowledgements

The authors would like to thank the canine patients and their families for their participation in this study in addition to the veterinary staff at the University of Wisconsin (UW) School of Veterinary Medicine for their excellent care and expertise including Rubi Hayim, CVT, Claire Quinn, DVM and Christina Lee, DVM. The authors also wish to thank the staff of the Program for Advanced Cell Therapy at the UW School of Medicine and Public Health for assisting with product release assays. Funding for this work was provided by the National Institutes of Health (NIH)/National Heart, Lung and Blood Institute T32 HL07899 (AW), NIH/National Center for Advancing Translational Sciences TL1 TR00375, SciMed Graduate Research Scholars fellowship (FS), the National Institute of General Medical Sciences/NIH T32 GM008692 and National Cancer Institute (NCI)/NIH T32 CA009135 (MMC), Czar’s Promise (DMV and CMC), St. Baldrick’s Foundation Empowering Immunotherapies for Childhood Cancer (EPICC) research grant, Lion’s Club International (CMC), and the Midwest Athletes Against Childhood Cancer (MACC) Fund (CMC). The Flow Cytometry core facility at the UW Carbone Cancer Center (UWCCC) is supported by a Cancer Center Support Grant from the NCI/NIH P30 CA014520. The contents of this article do not necessarily reflect the views or policies of the Department of Health and Human Services, nor does mention of trade names, commercial products, or organizations imply endorsement by the US Government.

