## Supplemental Figures for "Safety and Feasibility of Infusing *Ex Vivo* Expanded Allogeneic Canine Natural Killer Cells for the Treatment of Metastatic Solid Tumors"

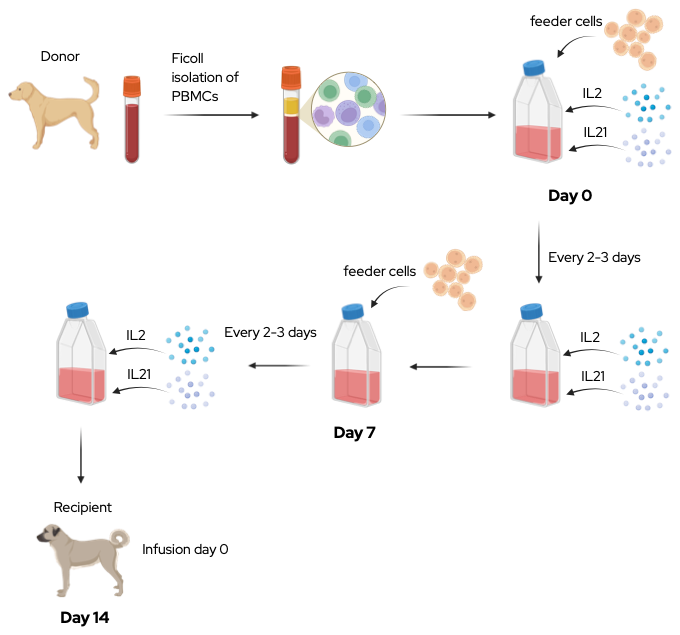

**Figure S1. Allogeneic NK cell expansion protocol.** PBMCs were isolated from peripheral blood of canine donors. IL-2 and IL-21 (as well as IL-12 in later experiments) were added every 2-3 days. Irradiated feeder cells were added on days 0 and 7 of the expansion

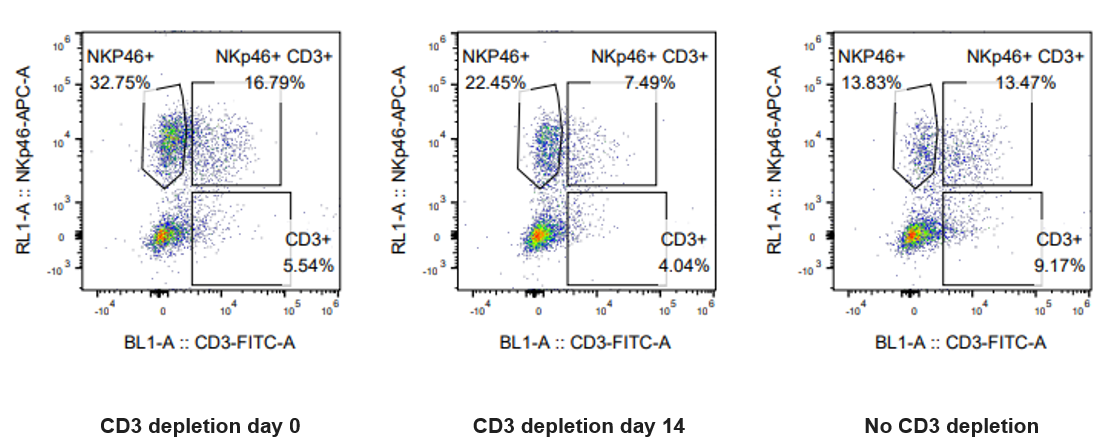

**Figure S2. Flow results of CD3 (T cell) depletion on day 0, 14, or not at all.** CD3 depletion experiments were performed on days 0 and 14 of expansion and compared to one another and to no depletion. All flow cytometry measurements were completed on day 15 of the expansions. CD3 depletion on day 0 yielded the largest percent NK cell expansion.

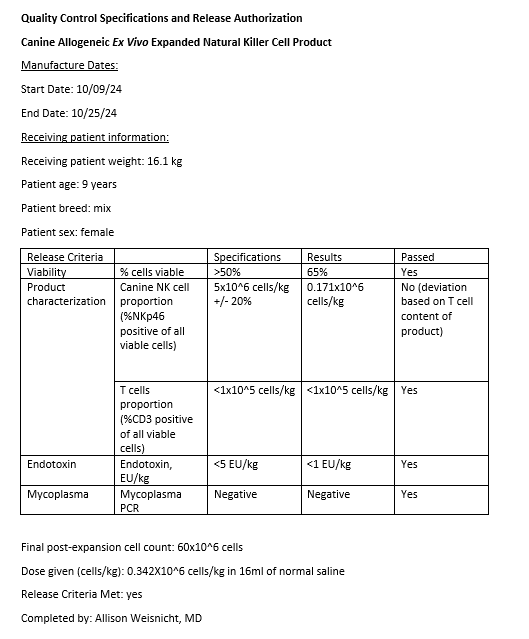

**Figure S3. PBMC-derived canine NK cell quality control release document.** The quality control release document includes percent viability, percent NK cells, percent T cells, endotoxin testing results, mycoplasma testing results, and cell dose.
